# mtDNA “Nomenclutter” and its Consequences on the Interpretation of Genetic Data

**DOI:** 10.1101/2023.11.19.567721

**Authors:** Vladimir Bajić, Vanessa Hava Schulmann, Katja Nowick

## Abstract

Population-based studies of human mitochondrial genetic diversity often require the classification of mitochondrial DNA (mtDNA) haplotypes into more than 5400 described haplogroups, and further grouping those into hierarchically higher haplogroups. Such secondary haplogroup groupings (e.g., “macro-haplogroups”) vary across studies, as they depend on the sample quality, technical factors of haplogroup calling, the aims of the study, and the researchers’ understanding of the mtDNA haplogroup nomenclature. Retention of historical nomenclature coupled with a growing number of newly described mtDNA lineages results in increasingly complex and inconsistent nomenclature that does not reflect phylogeny well. This “clutter” leaves room for grouping errors and inconsistencies across scientific publications, especially when the haplogroup names are used as a proxy for secondary groupings, and represents a source for scientific misinterpretation.

Here we explore the effects of phylogenetically insensitive secondary mtDNA haplogroup groupings, and the lack of standardized secondary haplogroup groupings on downstream analyses and interpretation of genetic data. We demonstrate that frequency-based analyses produce inconsistent results when different secondary mtDNA groupings are applied, and thus allow for vastly different interpretations of the same genetic data. The lack of guidelines and recommendations on how to choose appropriate secondary haplogroup groupings presents an issue for the interpretation of results, as well as their comparison and reproducibility across studies.

To reduce biases originating from arbitrarily defined secondary nomenclature-based groupings, we suggest that future updates of mtDNA phylogenies aimed for the use in mtDNA haplogroup nomenclature should also provide well-defined and standardized sets of phylogenetically meaningful algorithm-based secondary haplogroup groupings such as “macro-haplogroups”, “meso-haplogroups”, and “micro-haplogroups”. Ideally, each of the secondary haplogroup grouping levels should be informative about different human population history events. Those phylogenetically informative levels of haplogroup groupings can be easily defined using *TreeCluster*, and then implemented into haplogroup callers such as *HaploGrep3*. This would foster reproducibility across studies, provide a grouping standard for population-based studies, and reduce errors associated with haplogroup nomenclatures in future studies.

## Background

Mitochondrial DNA (mtDNA) is a circular non-recombining ∼16.5 kbp long extra-nuclear DNA. Exhibiting a tenfold higher mutation rate compared to nuclear DNA, with even higher mutation rates in the hypervariable segments (HVSI & HVSII), mtDNA shows great variation and diversity across individuals and can be used to study their evolutionary history. As mtDNA is inherited maternally, it is very informative about maternal population history and genetic structure, and it represents one of the commonly studied genetic markers [1, 2].

Early studies of human mtDNA diversity across the globe set the stage for the **mtDNA nomenclature** commonly used nowadays (for an overview see [3]). In 1992-1993, three milestone publications [4–6] on the mtDNA of Native Americans and indigenous peoples of Siberia not only proposed a genetic link between these two populations but also established a first nomenclature for mtDNA [3]. Additionally, they introduced the term ***haplogroup***, which stands for “haplotype group”, defined by the presence of specific mutations characteristic of a given branch of mtDNA phylogeny [3]. Due to these pioneering studies that introduced the nomenclature, the haplogroups A, B, C, and D were assigned to mtDNA types found in Native Americans and Siberians [6]. Later studies built upon this nomenclature, which turned out to be a suboptimal beginning of nomenclature given that humans originated in Africa and that the Americas were among the last regions humans reached. Over time, by researching diverse populations across the globe and accumulating mtDNA sequences, numerous “region-specific” haplogroups were identified [3]. Consistent with the original alphabetical nomenclature, newly discovered mtDNA haplogroups were assigned new letters. By the time the phylogenetically deepest splitting Sub-Saharan populations were studied, the haplogroups were designated “L” followed by a number [7] (Figure 1). Consequently, the current mtDNA nomenclature mostly reflects the history of research, and not necessarily the nested phylogenetic structure of the mtDNA tree.

**Figure 1.**
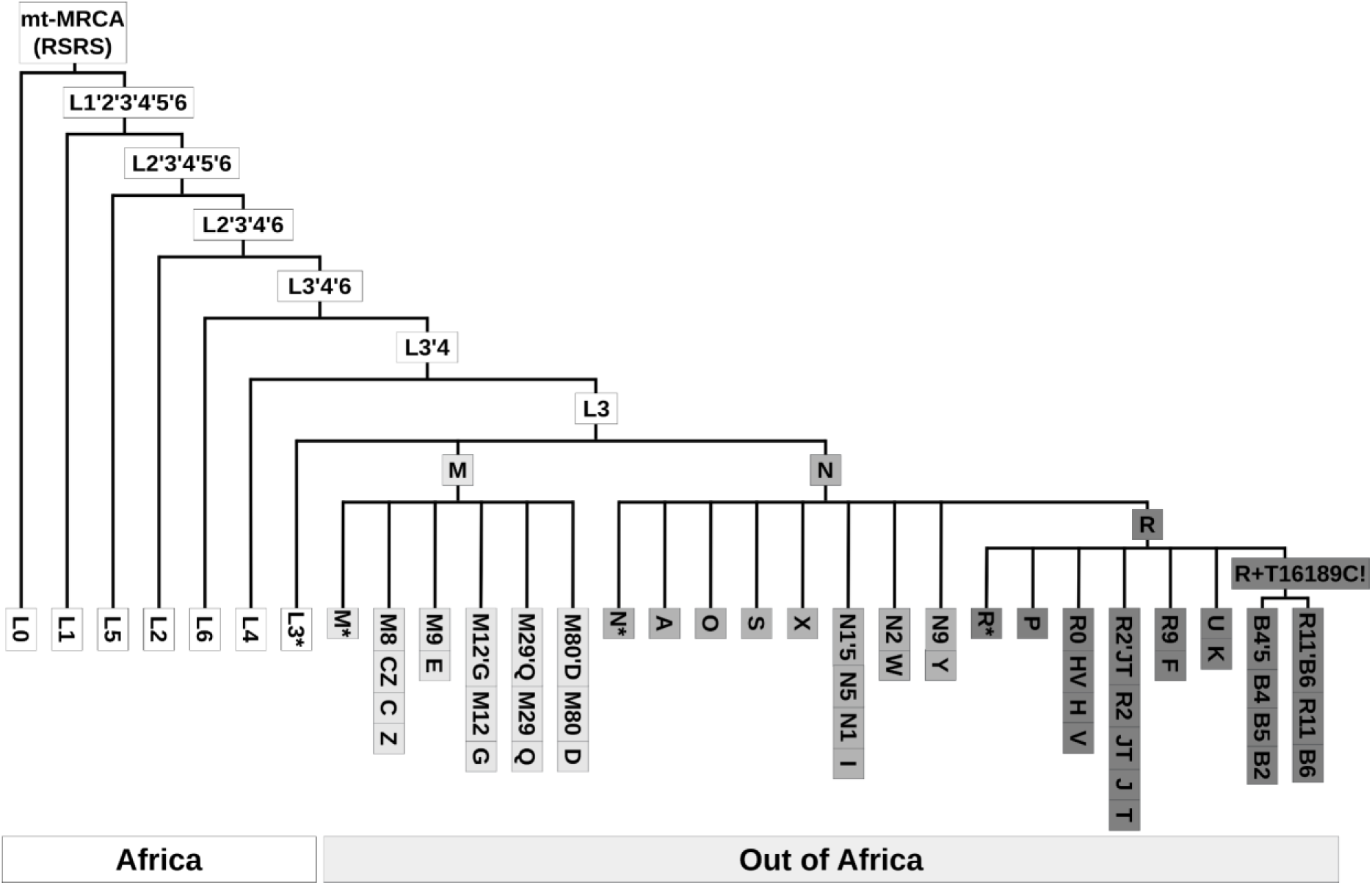
Schematic illustration of human mtDNA phylogeny from *PhyloTree build 17* (after van Oven 2015). The figure showcases the complexity in nomenclature as can be seen for example in how single capital letter haplogroups M, N, R, and A are nestled within two-character-long haplogroup L3.

To accommodate the ever-growing amounts of mtDNA sequences in the human mitochondrial tree, a systematic nomenclature was established (Text Box 1; for a detailed explanation see *“Cladistic notation for mitochondrial clusters”* in [8]). This nomenclature assumes that the first character of the haplogroup name is a capital letter, followed by a number and a lowercase letter in an alternating manner (e.g., C1d1), allowing easy naming for every new sub-branch. Consequently, the number of characters in a haplogroup name appears to reflect a phylogenetic level (which is not necessarily the case). Unfortunately, this naming convention became increasingly inconsistent with the discovery of new lineages and improvements in phylogeny.

##### Text Box 1 - Cladistic notation for mitochondrial clusters

Richards et al., 1998 outlined several principles for a cladistic notation for mitochondrial clusters:

**1)** Single capital letters denote principal clusters.

*E.g., A, B, C, D..*.

**2)** Nesting of clusters is permitted.

*E.g., C belongs to M, and M belongs to L3*.

**3)** Subclusters of single-capital-letter-coded clusters can have non-negative integer suffixes that follow hierarchical notation with alternating small letters and numbers.

*E.g., C1d1 is a subcluster of C1d which is a subcluster of C1 which belongs to C*.

**4)** Clade can be referred to using the names of its prominent subclades.

*E.g., CZ refers to the smallest monophyletic clade containing C and Z*.

**5)** Paraphyletic clusters can be temporarily labeled using an exponent suffix *.

*E.g., If U6b1, U6b2, and U6b3 are known then U6b* includes all U6b sequences not in already mentioned subclusters*.

The most comprehensive and commonly used resource for the organization and maintenance of the human mtDNA phylogeny is provided by **PhyloTree** [9, 10]. Between 2009 and 2016, PhyloTree periodically incorporated the ever-growing amount of mtDNA sequences into their phylogenetic tree, with the disclaimer that more branches are yet to be defined due to the biased sampling of populations.

For a given sample, **haplogroup calling** can be performed manually by checking if the haplogroup-specific mutations from *PhyloTree* are present or absent in the sample, or more commonly automatically using methods such as *HaploGrep3* [11]. ***HaploGrep3*** is a tool that assigns individual samples to the best-matching haplogroup according to the presence or absence of haplogroup-defining mutations from *PhyloTree*. The most recent *HaploGrep3* comes with several older *PhyloTree* versions, along with recent additions from forensics [12] and the ability to integrate external custom-made phylogenetic trees, which opens up the possibility for decentralized community-based updates.

Haplogroup callers such as *HaploGrep3* often output too many unique haplogroups to be useful for downstream population-based haplogroup frequency analyses. For example, *PhyloTree17-ForensicUpdate1.2* includes more than 6,300 haplogroups. This is why scientists often rely on custom secondary **Nomenclature-Based Groupings (NBG)** of haplogroups, in which they calculate the frequency of so-called **“macro-haplogroups”** or groups of related haplogroups (for example by summing frequencies of haplogroups B2a1a and B2a3 into macro-haplogroup B or even B2a). In contrast to the relatively well-defined concept of a **“haplogroup”**, other terms such as **“macro-haplogroup”**, **“super-haplogroup”**, **“major haplogroup”** and **“sub-haplogroup”** are not clearly defined. Those terms are instead often used to describe a relative relationship to the focal haplogroup, rather than representing clearly defined categories consistent across studies (e.g., one can say that B4a1 is a sub-haplogroup of haplogroup B4a, which belongs to macro-haplogroup B).

Two commonly occurring NBGs are **Single Character grouping (SC),** where grouping is based on the first haplogroup letter, and **Single Character and L with one digit grouping (SCL),** in which grouping is the same as SC except that haplogroups starting with “L” are grouped into more groups based on the first letter and the first digit (see Text Box 2 - mtDNA groupings). SCL makes more sense than SC as it better represents the haplogroup diversity within Africa. Besides those two NBGs, other modifications of secondary haplogroup groupings can be found in the literature. The newly released *Haplogrep3* introduces a slight modification to SCL by including HV as an additional category, resulting in 33 top-level haplogroups/clusters. In addition, custom groupings tailored to fit the study’s research question are also common in the literature (e.g., focusing on a particular haplogroup in more detail and lumping all the others in a single category). Consequently, the choice of NBG will influence the granularity (i.e. the phylogenetic level) of haplogroups and this can further influence the downstream analysis and interpretation of the data. If such grouping choices are uninformed by phylogeny, they can lead to creation of polyphyletic groups or pseudo-haplogroups, which may even corrupt study results [13].

##### Text Box 2 - mtDNA groupings

Throughout this paper, we consider three different types of *mtDNA groupings*:

**1) Haplogroup calling** - i.e. mtDNA haplogroup classification. A primary assignment of haplogroup names to haplotypes based on a set of mutations present in a given sequence. This step is commonly performed by haplogroup callers such as *HaploGrep3* [11], *HaploCart* [14], and *HaploGrouper* [15].

**2) Nomenclature-Based Groupings (NBG)** - i.e. creation of broader secondary mtDNA groupings based on traditional nomenclature, whose granularity level can vary from study to study (e.g., grouping samples assigned to haplogroups B4a1a1, B4a2b1a, B4a3, B4a4, B4a5 into B4a, or B4, or even B). Two commonly occurring NBGs are:

**i) Single Character grouping (SC)** - i.e. grouping based on the first haplogroup letter (e.g., grouping haplogroups C1b13c1 and C4a1a3a1 into C); or

**ii) Single Character and L with one digit grouping (SCL)** - i.e. grouping based on the first haplogroup letter except for those starting with “L”, for which the first letter and the first digit are used (e.g., grouping haplogroups L0d2c1a1 and L0k1a1c into L0).

**3) Algorithm-Based Groupings (ABG)** - i.e. computational grouping of mtDNA based on the sequence independent of traditional haplogroup nomenclature. Ideally, such groups of mtDNA should be in accordance with sequence similarities and/or mtDNA phylogeny. ABGs usually assign an arbitrary name or a number to a group of related mtDNA sequences. Such groupings can be obtained from mtDNA phylogenetic trees (e.g., using *TreeCluster* [16]), or directly from MSA in FASTA format (e.g., using *rhierBAPS* [17]).

The complexity of mitochondrial nomenclature and custom-made phylogenetic levels for secondary haplogroup groupings may lead to misunderstandings and inconsistencies between scientific publications. This is especially true when the lengths of haplogroup names are used as a proxy for their phylogenetic level and informativeness to create secondary groupings.

As mentioned above, the *PhyloTree17-ForensicUpdate1.2* has 6380 haplogroups, but nearly one-quarter of them have names that do not follow strict cladistic nomenclature, i.e. haplogroup names start with capital letters and are followed with alternating numbers and small letters (Table S1). For example, 1.8% of haplogroup names include two consecutive uppercase letters denoting nested haplogroups (e.g., haplogroups J and T are within JT; see Figure 1), 2.8% contain two consecutive lowercase letters (e.g., H3ag1), nearly 19% have one or more of the symbols *()@+” denoting mutated positions and/or lack of them (e.g., D5a2a1+@16172). Furthermore, many haplogroups were named by merging the names of two sub-branches, separated with an apostrophe (e.g., haplogroups M9a and M9b are nested within M9a’b). Besides the name itself, the phylogenetic relationship between haplogroups presents another obstacle. For example, haplogroups D1, D2, and D3 are nested within D4 instead of a more logical assumption of them being at the same or similar phylogenetic level. All these historically accumulated naming conventions make modern mtDNA nomenclature complex [3], unintuitive to navigate, and difficult to work with in bioinformatics pipelines, which is especially true for researchers new to the field.

In addition, the richness of defined haplogroup names depends on the studied populations, which results in systematic biases. European individuals and populations are often overrepresented in genetic databases [18]. Consequently, an mtDNA sample is more likely to be assigned to a haplogroup with a high resolution if they resemble well-studied and described populations, like Europeans. Moreover, *PhyloTree* has not been updated since 2016, which opens up the question about the future maintenance and standardization of mtDNA haplogroup nomenclature.

Many studies assume that **SC** groups are “macro-haplogroups” (e.g., assuming that both haplogroup L1 and L0d belong to “macro-haplogroup L”) (e.g., [19–22]). Such practice creates confusion as there is no haplogroup L, and since all non-L haplogroups are nested within L3. In reality, the hierarchically highest “L” haplogroup nomenclature designations all have two characters (L0 - L7) [10, 23]. Phylogenetically speaking, all haplogroups that contain “L” at the beginning of their name refer to the most recent common ancestor (MRCA) and should be grouped under mt-MRCA (that splits into L0 and L’1’2’3’4’5’6). What authors actually assume with SC grouping is a non-monophyletic grouping of haplogroups, whose names start with the letter L. Subsequently, this grouping leads to a disproportionate loss of phylogenetic resolution for the deepest lineages of the mtDNA phylogeny and thus has the strongest effect on African populations that harbor the highest frequencies and diversity of such haplogroups [24]. The effect on African populations is less severe when **SCL** is employed as it includes at least two characters for haplogroups whose name starts with the letter “L” (e.g., [25–27]. Even though slightly better than SCL, the 33 top-level haplogroups/clusters provided by *Haplogrep3* still do not account completely for the nested phylogenetic structure of haplogroups. For example, frequencies of M, N, and R are calculated separately as if R is not nested within N [22, 28]. Furthermore, there are small inconsistencies between *Haplogrep3* clusters depending on their phylogenies. For example, *Haplogrep3* “Cluster R” is defined differently, depending on whether PhyloTree 17 - Forensic Update 1.2 (https://haplogrep.i-med.ac.at/phylogenies/phylotree-furcrs@1.2/clusters/R; accessed on 30.09.2023) or PhyloTree 17.1 (https://haplogrep.i-med.ac.at/phylogenies/phylotree-rsrs@17.1/clusters/R; accessed on 30.09.2023) is used. In the first case “Cluster R” includes 198 haplogroups (including HV*, JT, and all that start with the letter R), while in the second case, it includes only the 39 haplogroups that start with R0 and all other haplogroups whose name starts with “R” are included in “Cluster N”.

In summary, complex nomenclature and lack of standardized or recommended grouping levels for haplogroups may introduce biases into downstream analyses and consequent interpretations of data. A better alternative to traditional nomenclature-based haplogroup groupings could be an **Algorithm-Based Grouping (ABG)** utilizing sequence similarities or phylogenetic trees. Such grouping could circumvent complex nomenclature and reduce human bias in choosing secondary haplogroup groupings.

Here we aim to explore how the categorization of human mitochondrial diversity into secondary NBGs can affect downstream haplogroup frequency-based analyses and our interpretations of the underlying genetic data. To do so, common NBGs are compared with ABGs produced by *rhierBAPS* and *TreeCluster*. Lastly, we propose that defining a standardized set of “macro-haplogroups”, “meso-haplogroups”, and “micro-haplogroups” in a phylogenetically meaningful way would be beneficial to the fields of population genetics, forensics, and medical genetics.

## Materials and methods

### Data and data preparation

We retrieved full mitochondrial sequences from 660 individuals free of relatives belonging to seven African Ancestry populations (ACB - African Caribbean in Barbados; ASW - African Ancestry in SW USA; MSL - Mende in Sierra Leone; GWD - Gambian in Western Division - Mandinka; ESN - Esan in Nigeria; YRI - Yoruba in Ibadan, Nigeria; LWK - Luhya in Webuye, Kenya) from the Phase 3 release of the 1KGP (1000 Genomes Project) [29]. The revised Cambridge Reference Sequence (rCRS) [30] was added as a reference for multiple sequence alignment. In addition, we added the Reconstructed Sapiens Reference Sequence (RSRS) [31] for rooting the phylogenetic tree. Positions known to interfere with multiple-sequence alignment were removed or altered manually using *Unipro UGENE v42.0* [32]. The spacer in the rCRS at position 3107 was changed from an “N” to an indel. The same was done for the spacers in the RSRS [31] at positions 523-524. The positions in the poly-c region 303–315 and 16183–16194 were removed from all sequences.

### Multiple sequence alignment (MSA)

MSA was performed using *MAFFT* [33]. Following good practice recommendations, manual post-alignment base correction in the region around the original 3107 rCRS spacer was performed using *Unipro UGENE*, ensuring that all indels align in the same pattern.

### Phylogeny construction and visualization

mtDNA phylogenetic trees were constructed by Maximum Likelihood (ML) with *IQ-TREE* [34]. The *ModelFinder Plus* setting was used to determine the best-fitting model by calculating the Bayesian Information Criterion (BIC) and choosing the model which minimizes the BIC. Phylogenetic trees were rooted using the RSRS [31]. To obtain a schematic reduced phylogenetic tree (Figure 3) results from secondary nomenclature-based groupings (NBG) and algorithm-based groupings (ABG) were combined to identify a minimal set of representative samples that capture all unique combinations of different groupings across individuals. The phylogenetic tree was visualized using the R package *ggtree* [35].

### mtDNA groupings

**Haplogroup calling** was performed using *Haplogrep3 (v.3.2.1)* command line tool [11], with FASTA as the input file, and the *PhyloTree17 - Forensic Update (rCRS Human mtDNA) Version 1.2 (phylotree-fu-rcrs@1.2) as the tree*. Based on the output of *HaploGrep3* we created two commonly occurring **secondary NBGs: SC** and **SCL** in R [36]. To obtain mitochondrial groupings based on mtDNA sequence similarity independent of traditional nomenclature, we performed two **ABGs:** rhierBAPS and TreeCluster.

***The rhierBAPS groupings (rhb)*** were performed directly on MSA FASTA using the R package *rhierBAPS* [17]. The *rhierBAPS* analysis was run with *keep.singletons=TRUE, max.depth=10,* and the default value for *n.pop*. For visualization purposes, we focused on the first 3 levels of the *rhierBAPS* output.

**The *TreeCluster* groupings (tc)** were performed on the mtDNA consensus trees outputted by *IQ-TREE* (chosen for its efficient computing times and likelihood maximization, and for its integrated model selection) using the command line program *TreeCluster* [16] with eight different threshold values (*-t*) ranging from 0.001 to 0.008, and method (*-m*) set to its default: *max_clade*. The Max Clade method of *TreeCluster* clusters the leaves of the provided phylogenetic tree ensuring that the maximum pairwise distance between leaves in the cluster is at most equal to the specified threshold. For visualization purposes, we focused on the output of *TreeCluster* with threshold values 0.003-0.006.

### Frequency-based analysis

#### Frequency bar plots

Frequency bar plots based on different mitochondrial groupings were created in R using packages *tidyverse* [37] and *ggpubr* [38].

#### Correspondence analysis (CA)

CA was performed using the R package *FactoMineR* [39] and visualized using the R package *factoextra* [40].

### Pairwise distance-based analyses

Pairwise distances between individuals from the same mitochondrial group were calculated in R using the packages *ape* [41] and *tidyverse* [37].

#### Multidimensional scaling (MDS)

MDS analysis was performed on the mtDNA pairwise distance matrix using the function *cmdscale()* from the R package *stats* [36] *and visualized using ggplot2* [38].

### Geographic map

The geographic map with populations was created using R packages *rworldmap* [42] *and tidyverse* [37].

### Literature meta-analysis

For a systematic overview of NBGs used in the literature, we assembled a list of research articles published in years 2020-2023 through Nature Publishing Group that contained the search term: *“africa* haplogrep* human* mtdna* haplogroup* OR africa* haplogrep* human* mitochondria* haplogroup*”* (Table S2). Each research article we categorized based on the NBG employed.

## Results and Discussion

### Comparison of NBG with ABG

To illustrate the differences between NBGs and ABGs we chose seven African Ancestry populations (AFR) from the 1000 Genomes Project (1KGP) Phase 3 (Figure 2A). Those populations are suitable as they harbor a high mtDNA diversity (Figure 2B, Figure S1, Figure S2) and many haplogroups, whose names are starting with the letter L that are often lumped together into “L macro-haplogroup” (assuming SC grouping). The majority of “non-L haplogroups” (A, B, C, D, H, J) in those populations are found in low frequency only in two populations from Americas (ACB - African Caribbean and ASW - African Ancestry SW) known to be admixed with non-African populations.

**Figure 2.**
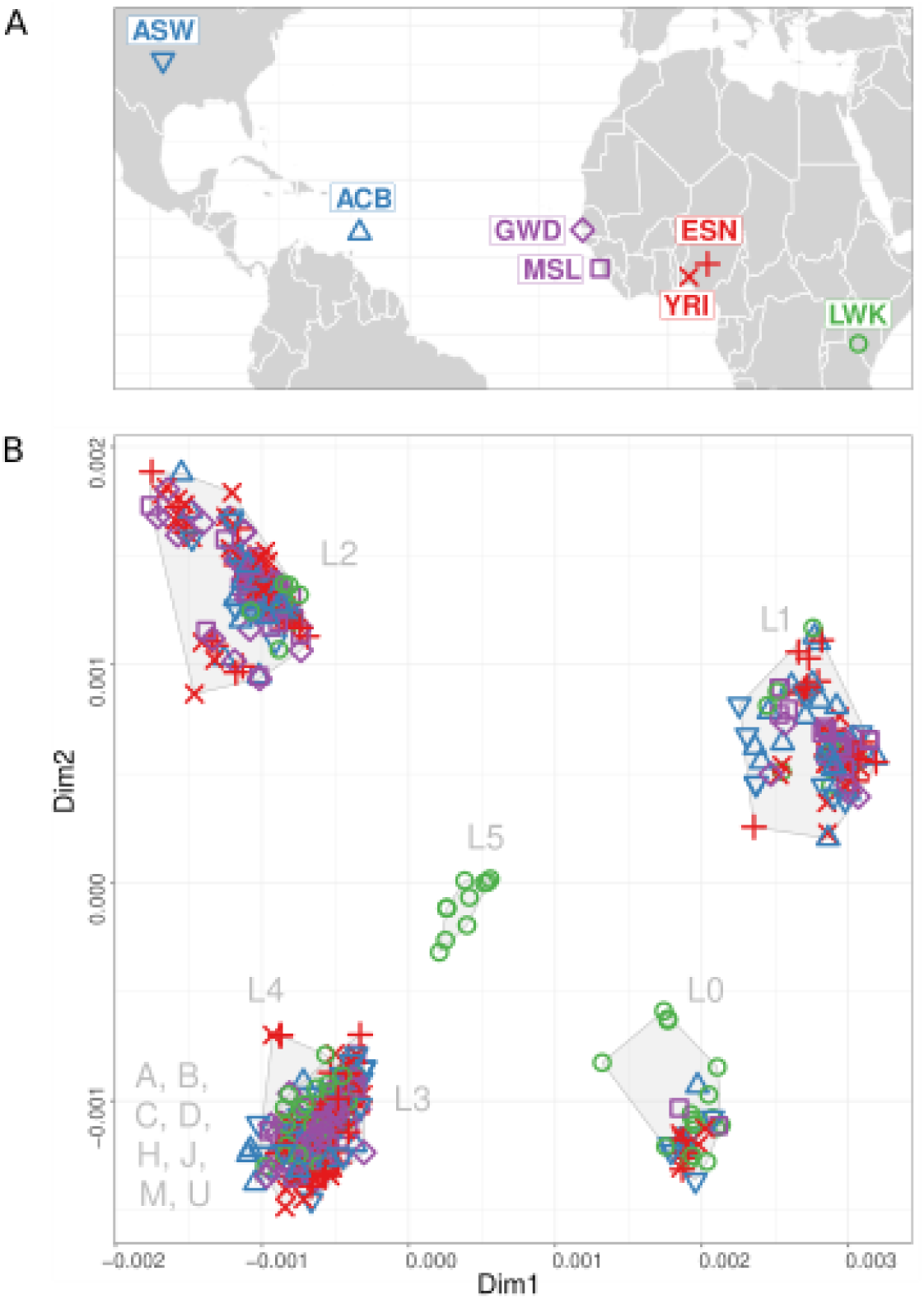
African Ancestry populations from the 1KGP Phase 3 and mtDNA pairwise-distance-based MDS plots. **A)** Map of sampling locations. ACB - African Caribbean in Barbados; ASW - African Ancestry in SW USA; MSL - Mende in Sierra Leone; GWD - Gambian in Western Division - Mandinka; ESN - Esan in Nigeria; YRI - Yoruba in Ibadan, Nigeria; LWK - Luhya in Webuye, Kenya. **B)** Two-dimensional MDS plots based on mtDNA pairwise distances colored according to populations. Population symbols correspond to those in panel A. SCL groupings are indicated with gray shadings and labels. Note that L3, L4, and “non-L” haplogroups are overlapping on this two-dimensional MDS plot.

The NBGs (SC and SCL) reflect phylogenetic structure poorly (Figure 3; for the full tree colored according to SCL see Figure S1). This is especially true for SC, which lumps the majority of mtDNA diversity into a single haplogroup and gives disproportionately more resolution to lineages commonly found outside of Africa. A noticeable improvement is seen with SCL, which gives slightly more resolution to haplogroups beginning with the letter L. On the other hand, *rhierBAPS* defines groups of sequences at several phylogenetic levels of resolution based on sequence similarity. Even though it provides more resolution for the phylogenetically deeper splitting haplogroups (i.e. haplogroups whose name starts with the letter L), *rhierBAPS* still produces phylogenetically inaccurate groups (e.g., phylogenetically related L2 haplogroups are not grouped together in *rhb_01*; a monophyletic group composed of L3b and L3d is nested within another group that would be monophyletic if it would include them in *rhb_02*).

**Figure 3:**
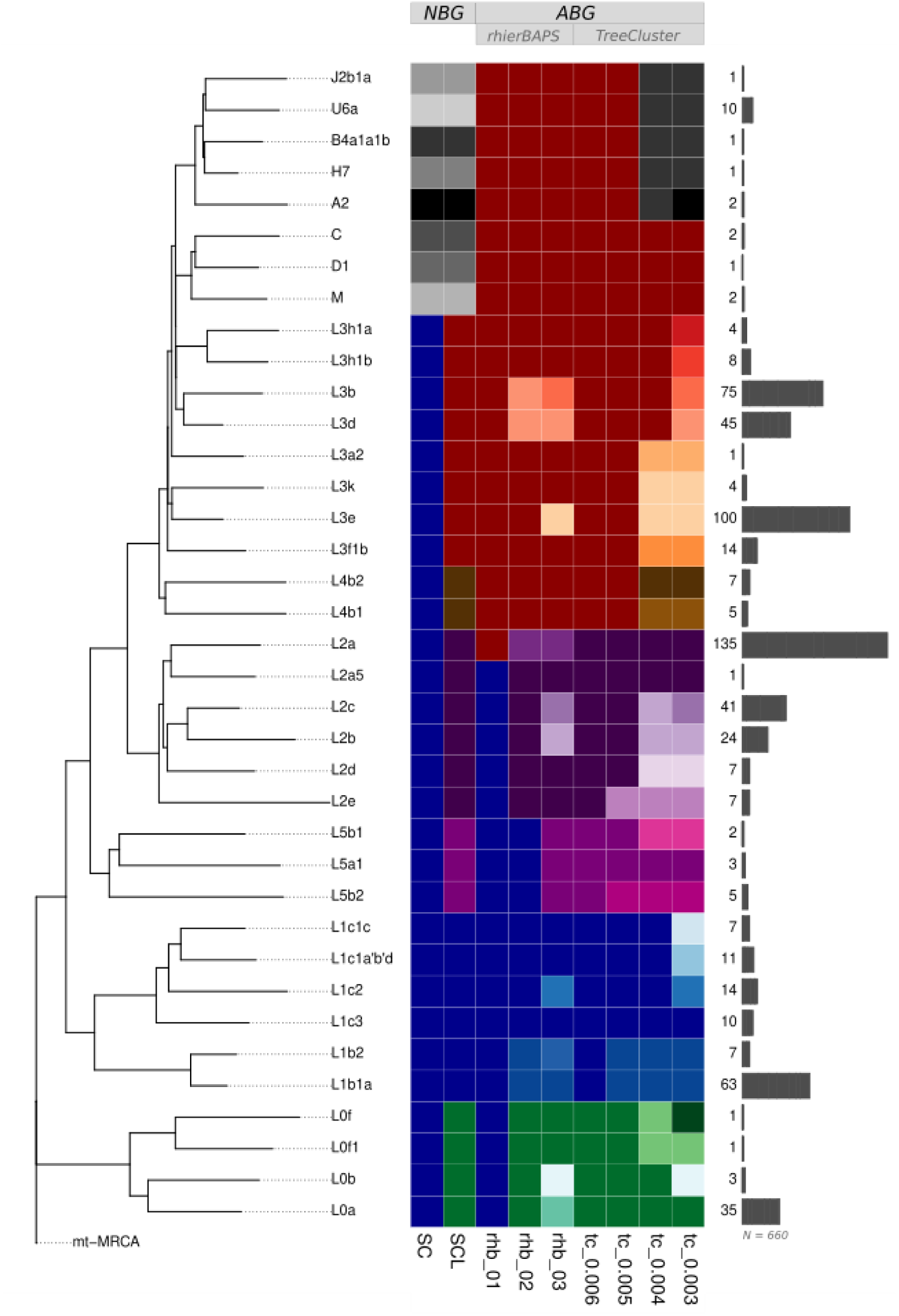
Comparison between nomenclature-based (NBG) and algorithm-based (ABG) groupings. The reduced phylogeny (left panel) is based on the minimal number of samples needed to capture the diversity based on combinations of all grouping designations. The total number of samples (N=660) belonging to each grouping is indicated with bar plots and numbers (right panel). Colors are used to visually distinguish groups within a grouping approach (vertically), meaning that each group is assigned a unique color. To illustrate the correspondence in grouping patterns between different grouping approaches (horizontally), similar colors and their gradients are used. See Table S3 for the results of NBGs and ABGs and their group names, which are here shown as different colors. Used abbreviations and the number of distinct groups produced by the grouping: SC - single-character grouping [9 groups]; SCL - single character and L with one digit grouping [14 groups]; rhb - rhierBAPS (level 1 [2 groups], level 2 [7 groups], and level 3 [16 groups]); tc - TreeCluster (threshold 0.006 [5 groups], threshold 0.005 [8 groups], threshold 0.004 [18 groups], and threshold 0.003 [29 groups]).

The best accordance with phylogeny is seen for *TreeCluster* groupings. This is expected as *TreeCluster* relies on a phylogenetic tree when making groupings, and the granularity of the groups can be adjusted with the threshold. In our case, the threshold of 0.006 (*tc_0.006*) resulted in the creation of groups equivalent to SCL groups L0, L1, L2, and L5. However, all other SCL groups are contained in the same group in *tc_0.006*.

As expected, decreasing the threshold values for the maximum pairwise distance between leaves in the cluster (*tc_0.005, tc_0.004, tc_0.003*) resulted in the creation of new groups predominantly in haplogroups whose name starts with the letter L (i.e. phylogenetically deeper splitting haplogroups). For example, only at *tc_0.003*, we can see the differentiation between non-L haplogroups, which are grouped into only 3 groups (first comprising haplogroup A, second comprising haplogroups B, J, U, and H, and third comprising haplogroups C, D, and M), while there are 26 groups whose haplogroup names start with letter L (which are all grouped into single L macro-haplogroup when SC is applied). Even though phylogenetically more meaningful and informative for African populations, such groupings would not be optimal for research focusing on non-African admixture in African Americans and African Caribbeans. However, this limitation could be resolved by further decreasing the threshold values, which would result in phylogenetically finer and more homogeneous groupings and thus distinction between non-L haplogroups. The choice of the exact threshold values would thus depend on the study and the question in mind.

### Mitochondrial grouping choices affect the interpretation of results

Different approaches to mitochondrial groupings are expected to have a direct effect on haplogroup frequency-based downstream analyses (e.g., haplogroup frequency plots such as pie charts or bar plots, and correspondence analysis). Here we demonstrate the effect of NBGs and ABGs on the outcome of such analysis. The SC approach distinguishes 9 groups from which the “L haplogroup” accounts for the majority of the data (Figure 4A). Besides being uninformative for African populations, SC could lead to a biologically misleading conclusion (e.g., that a given African population exhibits lower macro-haplogroup diversity when compared to non-African populations (see Table S2 for more examples).

**Figure 4.**
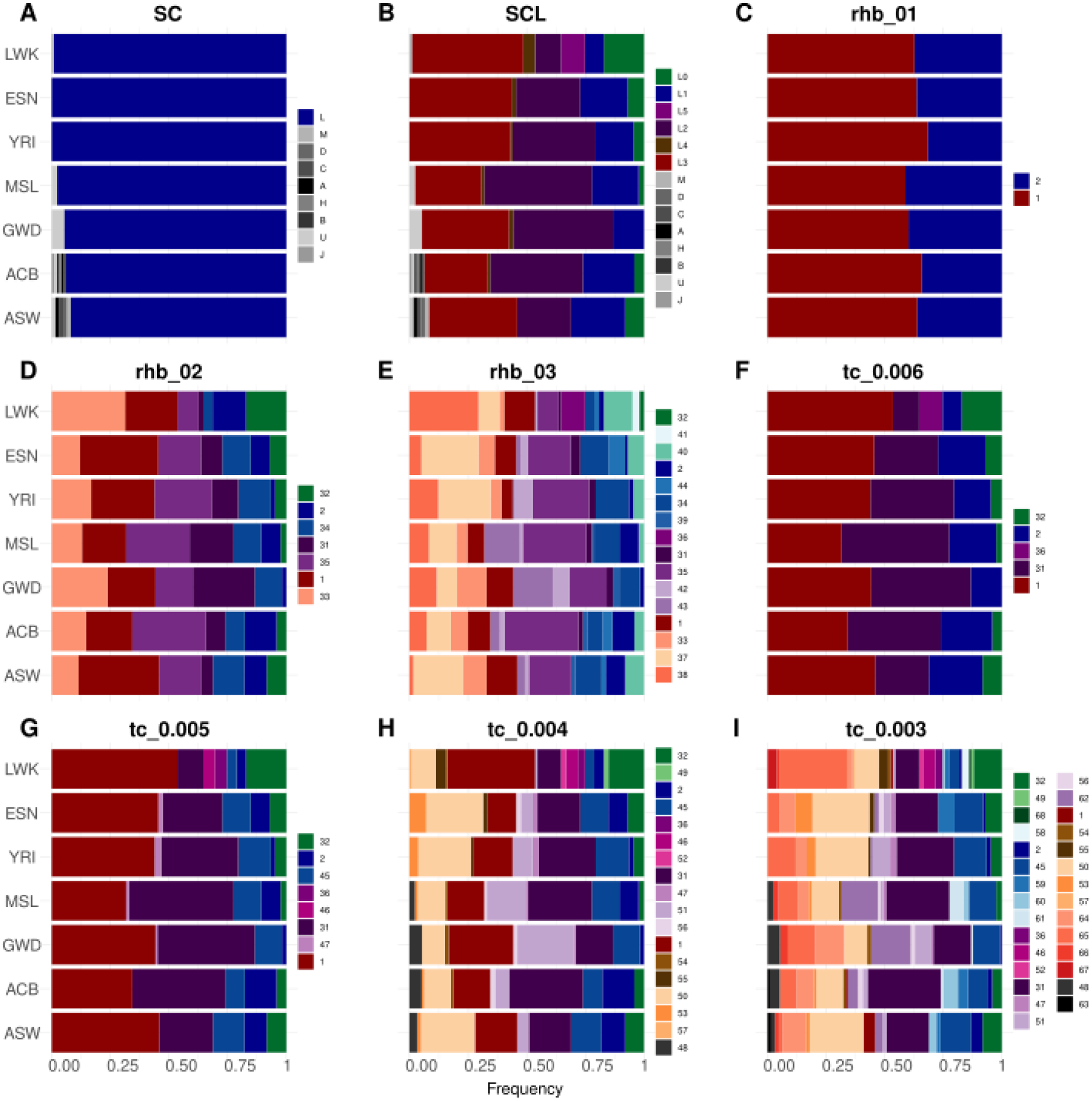
Frequency bar plots for African Ancestry populations from 1KGP based on mtDNA groups produced by NBGs: A) SC, and B) SCL; and ABGs: C-E) *rhierBAPS*, and F-I) *TreeCluster*. Note remarkable differences between SC and SCL as well as an increase in the number of haplogroups with increasing threshold values for ABGs. ACB - African Caribbean in Barbados; ASW - African Ancestry in SW USA; MSL - Mende in Sierra Leone; GWD - Gambian in Western Division - Mandinka; ESN - Esan in Nigeria; YRI - Yoruba in Ibadan, Nigeria; LWK - Luhya in Webuye, Kenya.

By implementing the SCL approach, the number of groups increases to 14 (Figure 4B), which starts to reveal the diversity of haplogroups in the sample. Comparisons between populations based on SCL allow more meaningful differentiation between African populations (e.g., L5 occurs only in LWK, while the majority of non-L haplogroups occur in two populations from the Americas). The comparison of frequency bar plots between SC and SCL is a striking example illustrating how NBGs could lead to drastically different interpretations of the same genetic data: The first level of *rhierBAPS* identified only two groups, the second level seven groups, and the third level 16 groups (Figure 4C-E). These different levels of granularity, however, might invite different conclusions. Based on *rhb_01,* there are hardly any differences between populations. Based on *rhb_02,* one could say that GWD shows the lowest haplogroup diversity, while all the other populations show comparable haplogroup diversity. Based on *rhb_03* YRI and GWD have the lowest haplogroup diversities and LWK clearly has the highest haplogroup diversity.

Decreasing the threshold value in *TreeCluster* groupings leads to a drastic increase in the number of defined groups (e.g., threshold of 0.006 produces 5 groups, while the threshold of 0.003 produces 29 groups). Even though an increased number of groups allows more precise inferences in phylogenetically more recent times (e.g., detection of admixture from non-African sources in African American and African Caribbean populations) it hinders visualization and interpretation of the data.

To assess how different the sequences belonging to the same secondary haplogroup groupings are, we calculated the pairwise distances between individuals belonging to the same group (Figure 5C-D, Figure S2). When examining the pairwise distances for NBGs, it is noticeable that the SC grouping “L” encompasses a large range of pairwise distances (Figure 5A, Figure S2A). This diversity is separated into 6 groups (L0-L5) when SCL is applied, which results in generally reduced mean pairwise distances within groups (Figure 5B, Figure S2B). However, the SCL groups still exhibit considerably high levels of pairwise differences.

**Figure 5.**
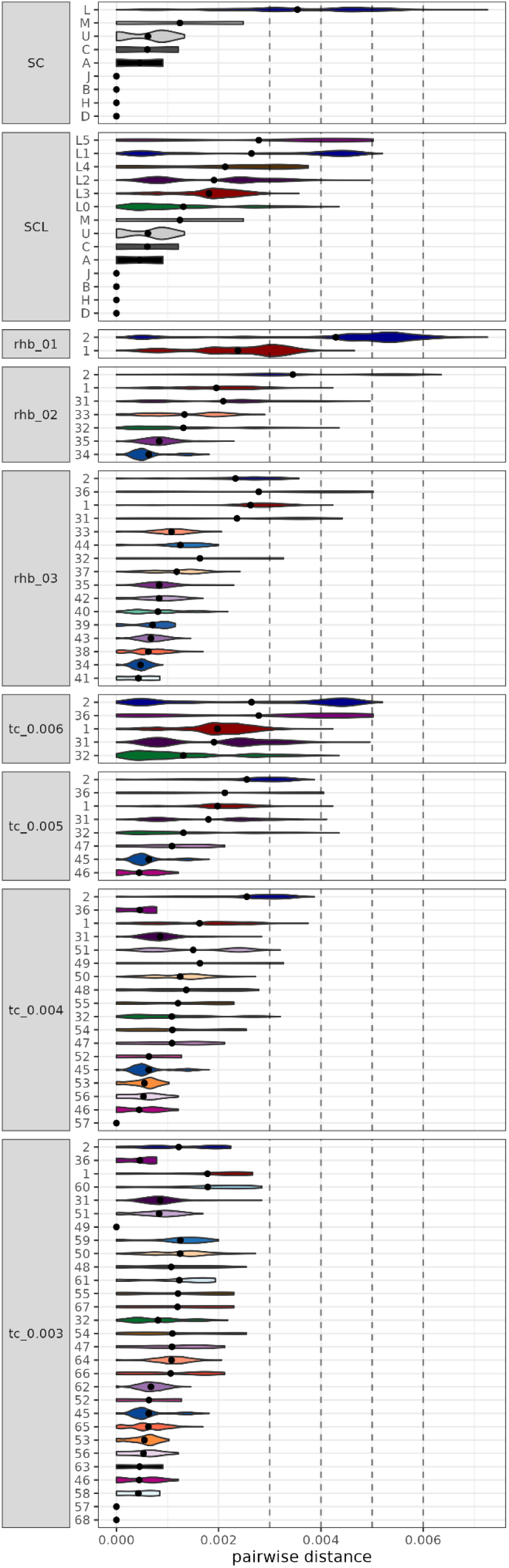
Violin plots of the mtDNA pairwise distances between individuals belonging to the same cluster produced by NBGs (SC and SCL), and ABGs (rhierBAPS and TreeCluster). Dashed lines indicate the threshold values used in TreeCluster runs. Note remarkable differences between SC and SCL, as well as an increase in the number of groupings with increasing rhierBAPS levels or decreasing threshold values for TreeCluster.

The ABGs allow us to define groups at different granularity levels by adjusting parameters and thresholds. The increased granularity from level 1 to level 3 in *rhierBAPS* groupings is reflected in lower within-group pairwise distances in higher levels (Figure 5C-E, Figure S2C-E). Still, level 3 (*rhb_03*) produces groups with very high variation in average and maximal within-group pairwise distances (e.g., groups 36, 31, and 1 show high (max values >0.004), while groups 34 and 41 show small within-group pairwise distances (max values <0.001)). On the other hand, *TreeCluster* allows strict control of the maximal within-group pairwise distances, and it uses information from a phylogenetic tree, which ultimately creates phylogenetically meaningful groups of comparable within-group pairwise distances, but at the expense of the increased number of groups (Figure 5F-I, Figure S2F-I).

#### Correspondence analysis

Correspondence Analysis (CA) is a multivariate statistical technique useful for visualizing the relative relationships between and within two groups of variables. In population genetic studies, CA is commonly used to explore the relationships among the frequencies of haplogroups and populations. Thus, CA is often used to estimate “similarity” or “difference” between a given set of populations based on their haplogroup frequencies. This, of course, heavily depends on the secondary haplogroup groupings (i.e., a phylogenetic level at which haplogroup calls are made). To investigate how different secondary haplogroup groupings affect CA, and ultimately the conclusions drawn from it, we compared eight different CA plots based on the NBGs (SC and SCL), and ABGs (*rhierBAPS* levels 2 and 3, and *TreeCluster* groupings with thresholds 0.006, 0.005, 0.004, and 0.003).

The SC and SCL CA plots exhibit striking differences in the clustering of populations on the first two dimensions. For the CA based on SC groupings (Figure 6A), the first two dimensions separate populations from Americas ASW and ACB from all the other populations that are almost indistinguishable from each other. The separation of populations from Americas is driven by the presence of non-African haplogroups (with the exception of haplogroup U).

**Figure 6:**
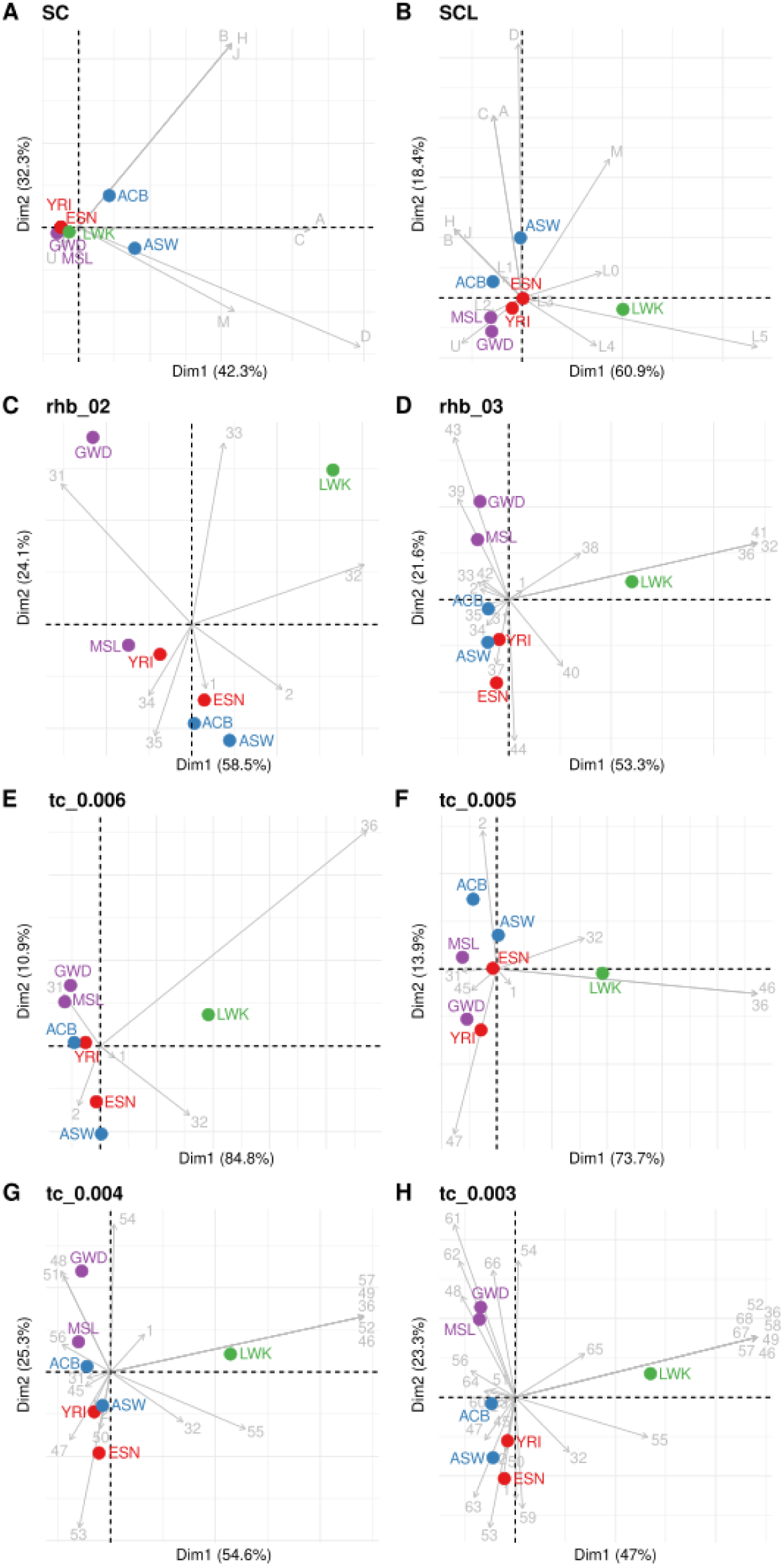
**Correspondence analysis (CA) plot of haplogroup frequencies of African Ancestry populations from 1KGP** based on groupings produced by A) SC, B) SCL, C) *rhierBAPS* level 2, D) *rhierBAPS* level 3, E) *TreeCluster* with threshold 0.006, F) *TreeCluster* with threshold 0.005, G) *TreeCluster* with threshold 0.004, H) *TreeCluster* with threshold 0.003. Arrows indicate the strength and direction of associations between mtDNA clusters and populations based on the first two dimensions (x- and y-axis). Colors indicate geographic regions: green - East Africa (LWK), blue - Americas (ACB and ASW), red - eastern West Africa (YRI and ESN), purple - western West Africa (GWD and MSL). ACB - African Caribbean in Barbados; ASW - African Ancestry in SW USA; MSL - Mende in Sierra Leone; GWD - Gambian in Western Division - Mandinka; ESN - Esan in Nigeria; YRI - Yoruba in Ibadan, Nigeria; LWK - Luhya in Webuye, Kenya.

The CA based on the SCL groupings (Figure 6B) still preserves the separation of populations from Americas ACB and ASW on the second dimension, while the first dimension separates East African LWK. This separation on the first dimension is primarily driven by haplogroup L5, which is unique to LWK in the dataset. The distinction between “L haplogroups” in SCL grouping thus adds valuable information for population differentiation within Africa.

Visually striking differences between SC and SCL CA could have a profound effect on the interpretation of the very same underlying genetic data. When interpreting the SC grouping, one can create a narrative, according to which the ACB and ASW populations are genetically more different from the continental African populations due to their non-African admixture. On the other hand, the SCL grouping could evoke an interpretation that the LWK population is genetically distinct from all of the other populations due to being East African and that the ACB and ASW populations outside of continental Africa are genetically distinct due to the presence of non-African haplogroups, yet still are genetically more similar to the West-African populations (ESN, GWD, MSL, YRI) than to the East-African populations (LWK). This could be explained by the trans-Atlantic slave trade, which mostly affected West-African populations [43]. While interpretations based on SC and SCL can both be inspired and supported (to some extent) by population history and thus both seem plausible, the question remains, which one of them is “better” or less biased for population genetics research?

Interpretations of CA based on ABGs pose a similar issue to NBGs. Even though East African LWK is always separated on the first axis, the separation of populations on the second axis is not always comparable across different ABGs (Figures 6C-H). For example, *rhierBAPS* Levels 2 (*rhb_02*) and 3 (*rhb_03*) show different clustering of populations, especially for GWD, which appears much more distant from MSI in *rhb_02* than in *rhb_03* (Figures 6C-D). The separation of populations from Americas is not obvious based on the first two dimensions in *TreeCluster* groupings except for *tc_0.005* (Figures 6E-H).

Based on comparisons between NBGs and ABGs we can conclude that the SC and SCL groupings put more emphasis on the non-L haplogroups due to the biases in haplogroup nomenclature, while the ABGs give more resolution to the phylogenetically diverse haplogroups whose names start with letter “L”.

However, these conclusions may be dependent on the dataset and haplogroup composition, as populations analyzed here predominantly consist of “L haplogroups”.

### Solutions to reduce biases in mitochondrial groupings

The complexity of traditional nomenclature and the biases it introduces in downstream analyses and interpretation of genetic data could be at least partially accounted for in several ways. Each requires different efforts for implementation and the likelihood of acceptance by the broader scientific community.

A radical solution to allow easier handling of mitochondrial haplogroups would be the development of a completely **new nomenclature** and the replacement of the current one. Even though several minor conflicts in mtDNA nomenclature for specific sets of haplogroups were already successfully resolved and incorporated into *PhyloTree* [10, 44–46], drastic changes in the mitochondrial nomenclature would come with issues and resistance in the scientific community, due to the widespread usage and well-established nature of the current nomenclature that ensures continuity with the previous mtDNA research [31, 47]. A possible new nomenclature system would require well-documented resources to aid with translation from an old to a new system, ensuring that previous research using old nomenclature could be related to future research and new nomenclature. Lastly, such a drastic change in nomenclature would have to be centralized, approved, and accepted by the global scientific community. Such centralization would also entail regular updating of the database and nomenclature to incorporate new mtDNA sequences, improving the accuracy and quality of haplogroup calling.

Alternatively, **algorithm-based groupings (ABGs)** could be useful in minimizing biases introduced by scientists when making secondary haplogroup groupings. For example, tools that group sequences based on sequence similarity [48, 49], or tools that make groups based on phylogenetic trees [16] can be very useful. Based on the results presented here, we suggest using *TreeCluster*, a tool that requires a phylogenetic tree as input, and can output different levels of secondary haplogroup groupings in seconds, even for phylogenies containing thousands of samples. Some ABGs can be labor-intensive, complicated to install and run, and might require additional data not readily available (e.g., rhierBAPS requires a multiple sequence alignment in FASTA format, and *TreeCluster* requires a phylogenetic tree in Newick format). Even though ABGs may introduce more reliable and reproducible groupings that are in better accordance with phylogeny, they are still not completely unbiased and still depend on arbitrarily chosen cutoffs. Lastly, running them independently for each dataset would reduce reproducibility and comparability across studies.

The most optimal solution, in our opinion, would be the **implementation of informed grouping recommendations based on ABGs**. Ideally, the definition of secondary haplogroup groupings should be performed **only once on the full mtDNA phylogeny, which is used as the basis for mtDNA nomenclature**. Currently, the most widely used nomenclatures are based on *PhyloTree* for which a phylogenetic tree in Newick format is not available. This is why we were not able to propose the actual set of standardized “macro-haplogroups”, “meso-haplogroups”, and “micro-haplogroups” for the current version of *PhyloTree*. Instead, we only illustrated how future mtDNA phylogenies aimed for updates on mtDNA haplogroup nomenclature could be used in combination with *TreeCluster* results to define sets of standardized secondary haplogroup groupings. This means that only authors publishing updated mtDNA phylogeny and nomenclature need to run *TreeCluster* in order to define secondary haplogroup groupings. Thus, no other authors would need to run *TreeCluster* themselves, which remains an option until the actual secondary haplogroup grouping standardization becomes available. Once the updated phylogeny and proposed secondary haplogroup groupings are available, researchers would only need to look up a simple table where each haplogroup name would be linked to its secondary haplogroup grouping memberships. Ideally, such information would be directly included in the output of haplogroup callers, such as *HaploGrep3*. This would ensure that users of a given haplogroup caller (i.e., *HaploGrep3*) would directly have at their disposal haplogroup predictions, as well as macro-, meso-, and micro-haplogroup memberships of predicted haplogroups (as illustrated in Table 1).

**Table 1.** A hypothetical illustration of how *HaploGrep3* output could be updated to include information about predefined and standardized micro-, meso-, and macro-haplogroups, as well as the mtHg-address unique to each haplotype. *TreeCluster* results based on the full *PhyloTree* phylogeny used for mtDNA haplogroup nomenclature could be used to define standardized sets of micro-, meso-, and macro-haplogroups, as well as mtHg-address. Certain *TreeCluster* thresholds informative about human migrations and population histories could be used to determine secondary haplogroup groupings (here indicated with red, blue, and green colors). For illustration purposes, we focus on nine hypothetical *TreeCluster* levels (*TreeCluster* runs with thresholds from 0 to 0.008). In reality, one can decide on any number of thresholds to be included in the mtHg-address. For the mtHg-address that distinguishes each haplotype, one would need to incorporate a *TreeCluster* run with a threshold of 0. Note that the names of micro-, meso-, and macro-Hg are determined based on the MRCA-haplogroup of all haplogroups belonging to the same group for the appropriate *TreeCluster* threshold. In the above example, tc_0.004 is used to define micro-haplogroups. Both B4a1a1b and J2b1a belong to the same tc_0.004 group, and thus R is determined as the most recent parent haplogroup for both of them. Please note that this table is completely fictional, and its sole purpose is to illustrate the concept of mtHg-address, how to achieve it, and how it could be integrated into *HaploGrep3* output. As such, this table is not a proposal for actual standardization.

| Sample ID | Haplogroup (Hg) | Secondary Haplogroup Groupings based on TreeCluster |  |  | TreeCluster runs with different thresholds |  |  |  |  |  |  |  |  | mthg-address |
| --- | --- | --- | --- | --- | --- | --- | --- | --- | --- | --- | --- | --- | --- | --- |
|  |  | Macro-Hg | Meso-Hg | Micro-Hg | tc_0.008 | tc_0.007 | tc_0.006 | tc_0.005 | tc_0.004 | tc_0.003 | tc_0.002 | tc_0.001 | tc_0.000 |  |
| S1 | L0f1 | L0a'b'f'g | L0f | L0f1 | 1 | 1 | 1 | 1 | 1 | 1 | 1 | 1 | 1 | 1.1.1.1.1.1.1.1.1 |
| S2 | L0a2a2a | L0a'b'f'g | L0a | L0a2 | 1 | 1 | 2 | 2 | 2 | 2 | 2 | 2 | 2 | 1.1.2.2.2.2.2.2.2 |
| S3 | L4b2a2b | L3'4 | L4 | L4b2 | 2 | 2 | 3 | 3 | 3 | 3 | 3 | 3 | 3 | 2.2.3.3.3.3.3.3.3 |
| S4 | B4a1a1b | L3'4 | N | R | 2 | 2 | 4 | 4 | 4 | 4 | 4 | 4 | 4 | 2.2.4.4.4.4.4.4.4 |
| S5 | J2b1a | L3'4 | N | R | 2 | 2 | 4 | 4 | 4 | 5 | 5 | 5 | 5 | 2.2.4.4.4.5.5.5.5 |
| S6 | A2n*2 | L3'4 | N | A | 2 | 2 | 4 | 4 | 6 | 6 | 6 | 6 | 6 | 2.2.4.4.6.6.6.6.6 |

**The optimal *TreeCluster* thresholds for macro-, meso-, and micro-haplogroups** should be carefully determined. The number of **macro-haplogroups** should ideally be under one hundred. They should be defined in such a way that many of the haplogroups often referred to as “macro-haplogroups” in the literature would also be included in *TreeCluster* defined macro-haplogroups. For example, on the level of macro-haplogroups, out-of-Africa lineages should be distinguishable from African lineages. The difficulty is that many deeply divergent mtDNA lineages that are uncommon might be assigned to their own macro-haplogroups. The usage of macro-haplogroups would be of greatest interest to researchers dealing with deep population histories or out-of-Africa migrations. **Meso-haplogroups** should ideally be able to distinguish lineages associated with major geographic areas or continents. Such, meso-haplogroups would be useful when investigating migrations and admixtures on global and cross-continental scales. And finally, **micro-haplogroups** should, at least to some extent, be capable of distinguishing lineages specific to some of the more recent major expansions in human history (e.g., agricultural expansions, expansions of language families, or some other prehistoric or historical events). Once the suitable thresholds are determined and applied, the group names produced by *TreeCluster* for each of those thresholds need to be translated into traditional nomenclature. This can be achieved by finding the haplogroup name of the MRCA-haplogroup of all haplogroups belonging to the same group for the given *TreeCluster* threshold (see example in Table 1).

In addition, we also propose the implementation of a supplementary mtDNA haplogroup nomenclature, the “**mtHg-address**”. Our inspiration for the mtHg-address derives from the **“SNP-address”** nomenclature [50], which was originally developed for pathogens. The **mtHg-address** could be defined using *TreeCluster* with several different threshold values (e.g., nine threshold values ranging from 0.008 to 0 as in Table 1). Each threshold produces monophyletic groups in which the maximum pairwise distance between samples is at most equal to the specified threshold. This nomenclature results in a unique name for each haplotype made of nine numbers separated by dots (here we use nine thresholds only for illustration purposes, but any other suitable number of thresholds could be selected). Each number in the name indicates the group membership at each of the predefined *TreeCluster* thresholds. For example, mtHg-address 2.2.4.4.4.5.5.5.5 and 2.2.4.4.6.6.6.6.6 indicate that two samples belong to the same monophyletic group based on *TreeCluster* thresholds from 0.008 to 0.005 but that they differ from each other starting with threshold 0.004. The mtHg-address thus provides a nomenclature that is meaningful regarding phylogeny, and it ensures that two identical haplotypes would have the same mtHg-address. As such, mtHg-address eliminates the possibility of pseudo-haplogroups or polyphyletic clades, which are often the result of historical haplogroup nomenclature, and the nested structure of haplogroups. As described before, some of the *TreeCluster* levels incorporated in the mtHg-address, carefully chosen based on their informativeness about mtDNA biogeography, would be used as guides for defining **macro-, meso-, and micro-haplogroups.** As suggested for standardization of micro-, meso-, and macro-haplogroups, the creation of mtHg-addresses should ideally be performed only once on the single mtDNA tree including all known mtDNA diversity, and then implemented in haplogroup callers such as *HaploGrep3*.

Lastly, similarly to mtHg-address inspired by pathogen research, rapid developments of tools capable of dealing with millions of sequences used in SARS-CoV-2 research could serve as an inspiration for future improvements of existing and development of new tools for mtDNA research [51–53]. For example, the mtDNA research community could benefit a lot if it could standardize the usage of the mutation-annotated tree (MAT) format for future mtDNA phylogeny and nomenclature updates. The MAT format efficiently encodes tree branch labels with parsimony-inferred mutations [51], and with slight adjustments for mtDNA research needs it could include mtHg-addresses and haplogroup labels. Moreover, minor updates of the evergrowing mtDNA phylogeny could be accomplished by developing tools similar to UShER (Ultrafast Sample placement on Existing tRees) [52], which would enable real-time updates of phylogenetic trees for mtDNA phylogeny without the need to construct entirely new phylogenetic trees. However, de-novo tree construction could still be performed less frequently (e.g., annually or biannually) for major updates of the mtDNA phylogeny.

The changes we proposed here would overcome problems related to traditional mtDNA nomenclature, reduce biases introduced by arbitrarily defined secondary nomenclature-based groupings, foster reproducibility across studies, and importantly, could be implemented into haplogroup callers such as *HaploGrep3*. It would also reduce the workload in any future studies, since the secondary haplogroup groupings and mtHg-address would already be pre-calculated and available to the community. Such changes would be of great importance and relevance for population genetics, forensics, and medical genetics.

## Declarations

## Data Availability

All scripts and output data are available on

GitHub: https://github.com/bajicv/mtDNA_nomenclutter

Zenodo: https://doi.org/10.5281/zenodo.10156923

All mentioned data, software, tools, and packages are publicly available and listed below:

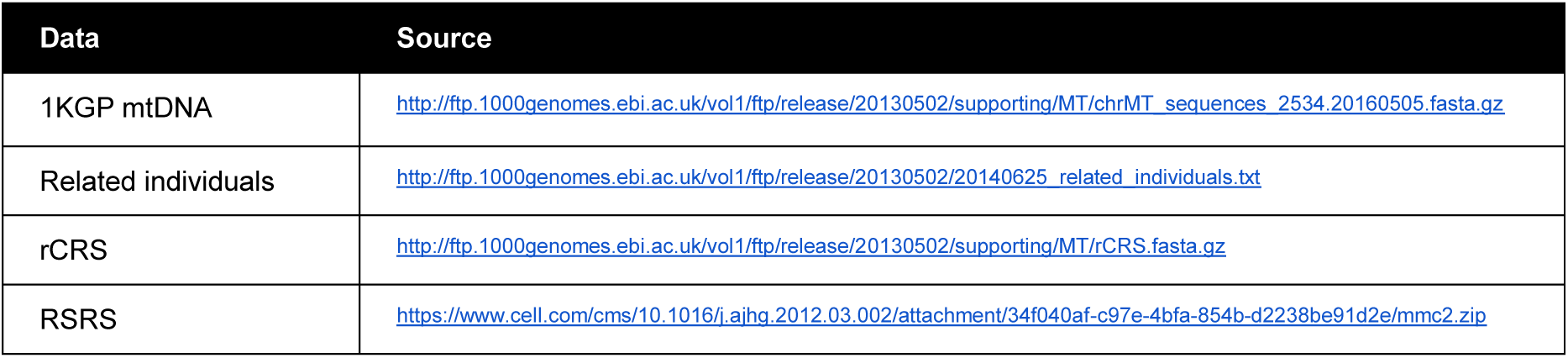

| Software/tool/package | Source |
| --- | --- |
| UGENE (v42.0) | http://ugene.net |
| MAFFT (v7.310) | http://mafft.cbrc.jp/alignment/software/ |
| IQ-TREE (v2.0.3 Linux) | http://www.iqtree.org |
| HaploGrep3 (v3.2.1) | https://haplogrep.i-med.ac.at/ |
| TreeCluster | https://github.com/niemasd/TreeCluster |
| R (v4.2.0) | https://www.cran.r-project.org/bin/macosx/ |
| R Studio (v1.1.383) | https://www.rstudio.com/products/rstudio/download/ |
| rhierBAPS | https://github.com/gtonkinhill/rhierbaps |
| ape | https://www.cran.r-project.org/web/packages/ape/index.html |
| seqinr | https://www.cran.r-project.org/web/packages/seqinr/index.html |
| phytools | https://www.cran.r-project.org/web/packages/phytools/index.html |
| ggtree | https://www.bioconductor.org/packages/release/bioc/html/ggtree.html |
| tidyverse | https://www.tidyverse.org |
| FactoMineR | https://www.cran.r-project.org/web/packages/FactoMineR/index.html |
| factoextra | https://www.cran.r-project.org/web/packages/factoextra/index.html |
| ggpubr | https://cran.r-project.org/web/packages/ggpubr/index.html |
| cowplot | https://cran.r-project.org/web/packages/cowplot/index.html |
| rworldmap | https://cran.r-project.org/web/packages/rworldmap/index.html |

## Authors’ contributions

Vladimir Bajić and Vanessa Hava Schulmann shared first authorship.

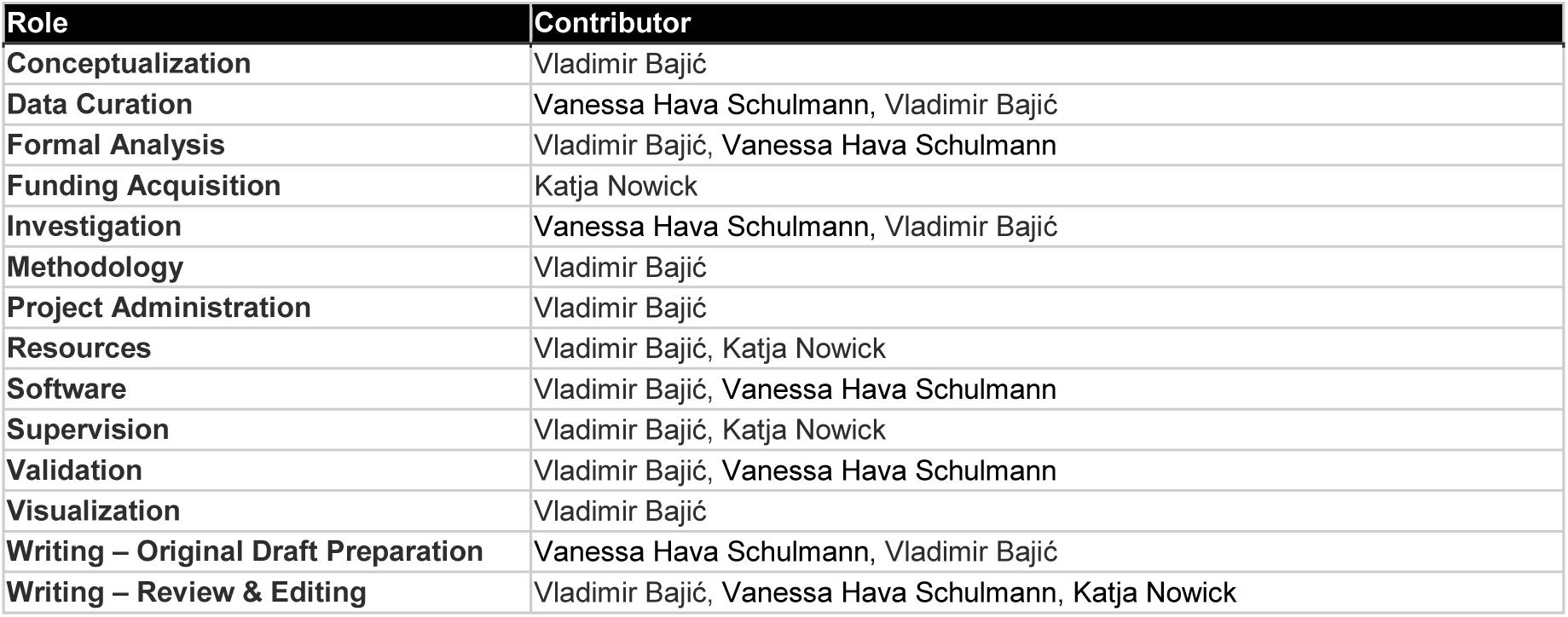

Authors' contributions Vladimir Bajić and Vanessa Hava Schulmann shared first authorship.
| Role | Contributor |
| --- | --- |
| Conceptualization | Vladimir Bajić |
| Data Curation | Vanessa Hava Schulmann, Vladimir Bajić |
| Formal Analysis | Vladimir Bajić, Vanessa Hava Schulmann |
| Funding Acquisition | Katja Nowick |
| Investigation | Vanessa Hava Schulmann, Vladimir Bajić |
| Methodology | Vladimir Bajić |
| Project Administration | Vladimir Bajić |
| Resources | Vladimir Bajić, Katja Nowick |
| Software | Vladimir Bajić, Vanessa Hava Schulmann |
| Supervision | Vladimir Bajić, Katja Nowick |
| Validation | Vladimir Bajić, Vanessa Hava Schulmann |
| Visualization | Vladimir Bajić |
| Writing – Original Draft Preparation | Vanessa Hava Schulmann, Vladimir Bajić |
| Writing – Review & Editing | Vladimir Bajić, Vanessa Hava Schulmann, Katja Nowick |

## Ethics approval and consent to participate

Not applicable.

## Consent for publication

Not applicable.

## Acknowledgments

A preprint version of this article has been peer-reviewed and recommended by PCIEvolBiol (https://doi.org/10.24072/pci.evolbiol.100716).

We thank all members of the Human Biology Group at FU-Berlin for fruitful discussions and comments, especially Melanie Sarfert for critically reading our manuscript and giving us constructive feedback. We also thank Chiara Barbieri and Sandra Oliveira for inspiring discussions and comments.

## Abbreviations

NBG: Nomenclature-Based Groupings
SC: Single Character grouping
SCL: Single Character and L with one digit grouping
ABG: Algorithm-Based Groupings
mtDNA: Mitochondrial DNA
rCRS: The revised Cambridge Reference Sequence
RSRS: The Reconstructed Sapiens Reference Sequence
MDS: Multidimensional scaling
CA: Correspondence analysis
*rhb_01*: Results of *rhierBAPS* level 1
*rhb_02*: Results of *rhierBAPS* level 2
*rhb_03*: Results of *rhierBAPS* level 3
*tc_0.006*: Results of *TreeCluster* with the threshold value of 0.006
*tc_0.005*: Results of *TreeCluster* with the threshold value of 0.005
*tc_0.004*: Results of *TreeCluster* with the threshold value of 0.004
*tc_0.003*: Results of *TreeCluster* with the threshold value of 0.003

## Population abbreviations

ACB: African Caribbean in Barbados
ASW: African Ancestry in SW USA
MSL: Mende in Sierra Leone
GWD: Gambian in Western Division – Mandinka
ESN: Esan in Nigeria
YRI: Yoruba in Ibadan, Nigeria
LWK: Luhya in Webuye, Kenya

## Supplementary Material

### Supplementary Tables

**Table S1.**
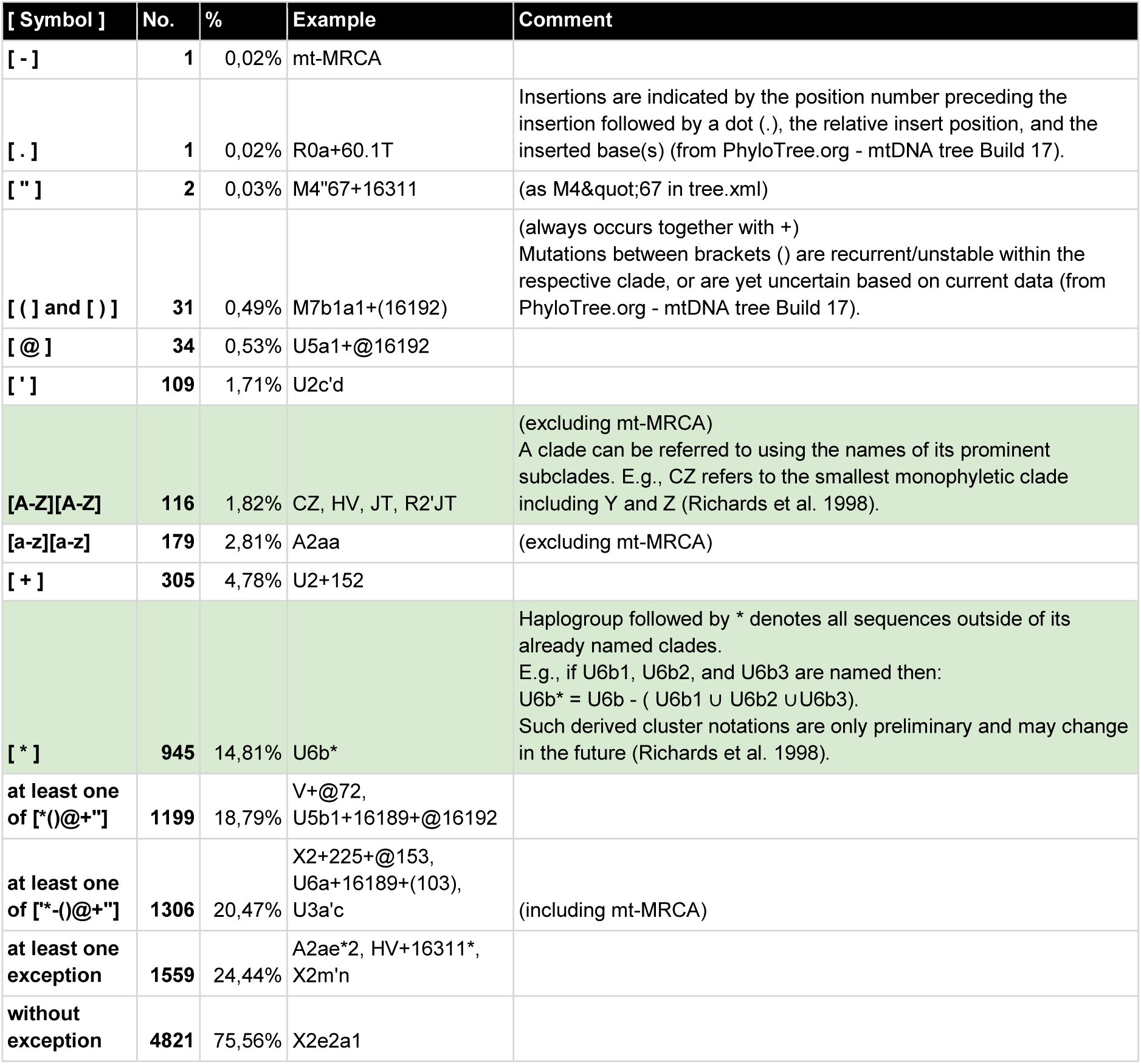
The number of haplogroup names from *phylotree-fu-rcrs-1.2* (total of 6380 haplogroups) that do not follow strict cladistic nomenclature with haplogroup names starting with capital letters followed with alternating numbers and small letters. Note that the “cladistic notation for mitochondrial clusters” established by Richards et al. 1998 (see Test Box 1) permits composed names to refer to monophyletic clades based on subclades (e.g., CZ), and usage of * that designates all sequences that do not belong to any of the already named subclades (green rows in the table).

**Table S2.** Overview of Nomenclature-Based Groupings (NBGs) used in the research articles published in years 2020-2023 through Nature Publishing Group (https://www.nature.com/search/advanced) with the search term: “africa* haplogrep* human* mtdna* haplogroup* OR africa* haplogrep* human* mitochondria* haplogroup*”. Each research article was categorized based on the NBG employed. Available at GitHub: https://github.com/bajicv/mtDNA_nomenclutter/blob/main/supplements/Table_S2_mtDNA_Nomenclutter.xlsx and at Zenodo: https://doi.org/10.5281/zenodo.10156923.

**Table S3.** Results of NBGs and ABGs used for plotting Figure 3. For exact correspondence between colors and names of groups see scripts/my_colors.R at https://zenodo.org/records/10156923

|  | n | NBG |  | ABG |  |  |  |  |  |  |
| --- | --- | --- | --- | --- | --- | --- | --- | --- | --- | --- |
|  |  | SC | SCL | rhierBAPS |  |  | TreeCluster |  |  |  |
|  |  |  |  | rhb_01 | rhb_02 | rhb_03 | tc_0.006 | tc_0.005 | tc_0.004 | tc_0.003 |
| J2b1a | 1 | J | J | 1 | 1 | 1 | 1 | 1 | 48 | 48 |
| U6a | 10 | U | U | 1 | 1 | 1 | 1 | 1 | 48 | 48 |
| B4a1a1b | 1 | B | B | 1 | 1 | 1 | 1 | 1 | 48 | 48 |
| H7 | 1 | H | H | 1 | 1 | 1 | 1 | 1 | 48 | 48 |
| A2 | 2 | A | A | 1 | 1 | 1 | 1 | 1 | 48 | 63 |
| C | 2 | C | C | 1 | 1 | 1 | 1 | 1 | 1 | 1 |
| D1 | 1 | D | D | 1 | 1 | 1 | 1 | 1 | 1 | 1 |
| M | 2 | M | M | 1 | 1 | 1 | 1 | 1 | 1 | 1 |
| L3h1a | 4 | L | L3 | 1 | 1 | 1 | 1 | 1 | 1 | 67 |
| L3h1b | 8 | L | L3 | 1 | 1 | 1 | 1 | 1 | 1 | 66 |
| L3b | 75 | L | L3 | 1 | 33 | 38 | 1 | 1 | 1 | 65 |
| L3d | 45 | L | L3 | 1 | 33 | 33 | 1 | 1 | 1 | 64 |
| L3a2 | 1 | L | L3 | 1 | 1 | 1 | 1 | 1 | 57 | 57 |
| L3k | 4 | L | L3 | 1 | 1 | 1 | 1 | 1 | 50 | 50 |
| L3e | 100 | L | L3 | 1 | 1 | 37 | 1 | 1 | 50 | 50 |
| L3f1b | 14 | L | L3 | 1 | 1 | 1 | 1 | 1 | 53 | 53 |
| L4b2 | 7 | L | L4 | 1 | 1 | 1 | 1 | 1 | 55 | 55 |
| L4b1 | 5 | L | L4 | 1 | 1 | 1 | 1 | 1 | 54 | 54 |
| L2a | 135 | L | L2 | 1 | 35 | 35 | 31 | 31 | 31 | 31 |
| L2a5 | 1 | L | L2 | 2 | 31 | 31 | 31 | 31 | 31 | 31 |
| L2c | 41 | L | L2 | 2 | 31 | 43 | 31 | 31 | 51 | 62 |
| L2b | 24 | L | L2 | 2 | 31 | 42 | 31 | 31 | 51 | 51 |
| L2d | 7 | L | L2 | 2 | 31 | 31 | 31 | 31 | 56 | 56 |
| L2e | 7 | L | L2 | 2 | 31 | 31 | 31 | 47 | 47 | 47 |
| L5b1 | 2 | L | L5 | 2 | 2 | 36 | 36 | 36 | 52 | 52 |
| L5a1 | 3 | L | L5 | 2 | 2 | 36 | 36 | 36 | 36 | 36 |
| L5b2 | 5 | L | L5 | 2 | 2 | 36 | 36 | 46 | 46 | 46 |
| L1c1c | 7 | L | L1 | 2 | 2 | 2 | 2 | 2 | 2 | 61 |
| L1c1a'b'd | 11 | L | L1 | 2 | 2 | 2 | 2 | 2 | 2 | 60 |
| L1c2 | 14 | L | L1 | 2 | 2 | 44 | 2 | 2 | 2 | 59 |
| L1c3 | 10 | L | L1 | 2 | 2 | 2 | 2 | 2 | 2 | 2 |
| L1b2 | 7 | L | L1 | 2 | 34 | 39 | 2 | 45 | 45 | 45 |
| L1b1a | 63 | L | L1 | 2 | 34 | 34 | 2 | 45 | 45 | 45 |
| L0f | 1 | L | L0 | 2 | 32 | 32 | 32 | 32 | 49 | 68 |
| L0f1 | 1 | L | L0 | 2 | 32 | 32 | 32 | 32 | 49 | 49 |
| L0b | 3 | L | L0 | 2 | 32 | 41 | 32 | 32 | 32 | 58 |
| L0a | 35 | L | L0 | 2 | 32 | 40 | 32 | 32 | 32 | 32 |
|  | 660 | 9 | 14 | 2 | 7 | 16 | 5 | 8 | 18 | 29 |
|  | SUM | Number of groups |  |  |  |  |  |  |  |  |

### Supplementary Figures

**Figure S1.**
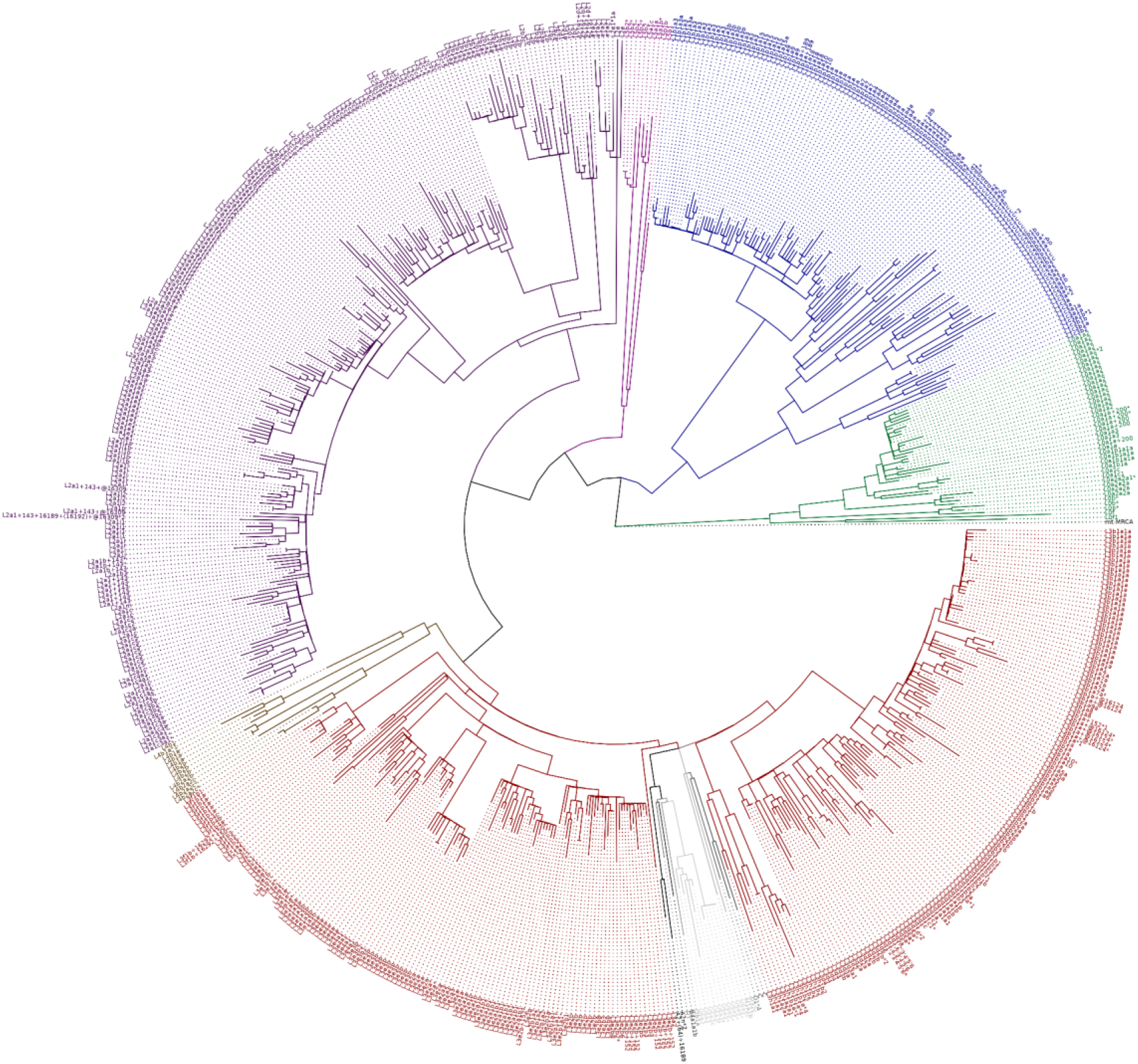
The mtDNA ML consensus trees of 660 samples belonging to seven African Ancestry populations (ACB, ASW, MSL, GWD, ESN, YRI, LWK) from the Phase 3 release of the 1KGP. Colors indicate SCL membership (single character and L with one-digit grouping). RSRS (mt-MRCA) was added for rooting the tree. The labels indicate haplogroup reported by *HaploGrep3*.

**Figure S2.**
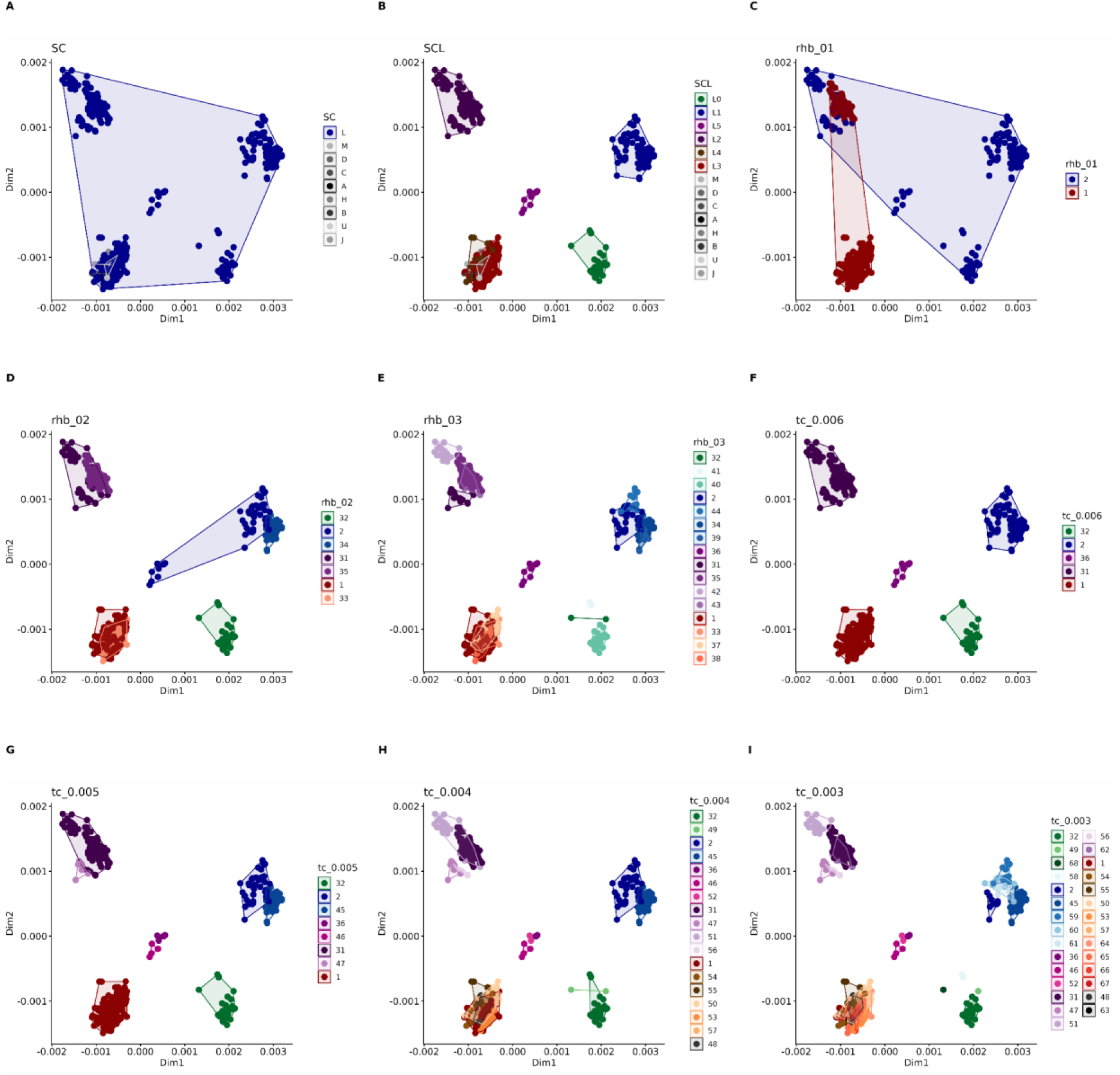
MDS plots based on the mtDNA pairwise distances between individuals colored by NBGs: A) SC, and B) SCL; and ABGs: C-E) *rhierBAPS*, and F-I) *TreeCluster*. Note remarkable differences between SC and SCL as well as an increase in the number of haplogroups with increasing threshold values for ABGs.

